# Investigating *Bacillus anthracis* genomic diversity and trait-specific lineages in an endemic area in northern Tanzania through a combination of traditional and culture-free sequencing approaches

**DOI:** 10.64898/2025.12.04.692280

**Authors:** Antonia Hilbig, Matej Medvecky, Tiziana Lembo, Blandina T. Mmbaga, Ireen Kiwelu, Deogratius Mshanga, Shabani K. Motto, Zacharia E. Makondo, Boaz Wadugu, Henri O. Arola, Simo Nikkari, Matthew P. Rubach, John A. Crump, Samantha J. Lycett, Roman Biek, Taya L. Forde

## Abstract

Anthrax, caused by *Bacillus anthracis* (BA), is a prominent neglected zoonosis with major impacts on human, livestock, and wildlife health. Despite this, limited genomic investigation at the One Health interface constrains current understanding of BA transmission and of the ecological and host factors shaping its diversity and population structure. This includes the possibility of host-specific BA lineages, given that anthrax outbreaks often disproportionally affect individual species. This study characterises the genomic diversity of BA in an endemic area, the Ngorongoro Conservation Area (NCA), in northern Tanzania. We analysed 213 BA genomes from livestock, wildlife and humans from cultured isolates combined with a culture-free targeted capture (TC) approach. NCA sequences formed a distinct genetic cluster compared with those from surrounding areas, and we observed surprisingly high levels of strain diversity within apparent epidemiological clusters, as well as within single animals, though strain diversity was lowest at the within host scale. We found limited evidence for seasonal clustering of cases as well as for BA lineages clustering by host species. This indicates that disproportional impacts on certain species during outbreaks are more likely driven by host ecology factors or, hypothetically, by accessory parts of the bacterial genome not represented in our data. TC-derived data significantly expanded the range of host species and geographic locations for genomic analysis, demonstrating the value of this approach. Although TC data may contain artefactual variation, shared SNP profiles between isolate- and TC-derived genomes gave confidence in its use for genotyping. Our analysis demonstrates unexpectedly high BA strain diversity and limited population structure in this endemic area across a range of spatial scales, including within-host. It further highlights the need for high density sampling and adaptable sequencing strategies to generate adequate BA genomic datasets that can enable informative molecular epidemiological studies of anthrax at the One Health interface.

**Author Summary:** Anthrax continues to threaten the health of people, livestock, and wildlife in many parts of the world, yet we still know surprisingly little about how this disease spreads in nature. One major gap is understanding why some species are affected more strongly during outbreaks, despite assumed equal susceptibility. To investigate this, we studied the genomic diversity of the anthrax-causing bacterium *Bacillus anthracis* in a large conservation area in northern Tanzania. We combined two ways of generating genetic data: traditional laboratory culture and a culture-free method that allowed us to recover bacterial DNA from a wider range of samples.

By analysing a uniquely large and species-diverse dataset of over 200 bacterial genomes from people, livestock and wildlife, we found that bacteria from the study area formed a clearly defined group compared to those from surrounding regions. We also discovered unexpectedly high diversity of strains, not only across the landscape but even within single animals. Despite this diversity, we saw little evidence that certain bacterial lineages are tied to specific host species. Our results suggest that the behaviour and ecology of different animals, rather than host-adapted lineages of the bacterium, likely explain why some species are more affected than others.

## Introduction

Neglected zoonotic diseases continue to have substantial global impacts on human and animal health and livelihoods, particularly in vulnerable populations in low- and middle-income countries, LMICs (1). Anthrax, caused by *Bacillus anthracis* (BA), is a prime example of these diseases (2). While anthrax outbreaks are rare in high-income countries, where overall morbidity and mortality are minimal (3, 4), in LMICs, particularly in remote rural communities, anthrax is endemic and remains a substantial public health challenge (5, 6). Here, frequent outbreaks cause considerable livestock and wildlife losses, as well as human illness and deaths (7–10).

The infectious form of the BA bacterium is a highly resilient spore that can retain infectivity for decades in the environment (3, 11–13). Spore uptake by herbivorous livestock and wildlife typically happens during grazing, when infectious material is ingested or inhaled from the soil or plant material (14). Infections will almost always end in sudden death of the animal with subsequent release of new infectious material from the carcass (15, 16). Human BA infection occurs most frequently during the handling or consumption of the meat or hide of animals that have died from the disease, leading to cutaneous lesions, or gastrointestinal or respiratory infections. Cutaneous lesions tend to be self-limiting, although may lead to blood stream infection, whereas gastrointestinal disease is often severe and respiratory infections are typically fatal (17–19).

Although BA is known for low genomic variation, also referred to as monomorphism (20, 21), whole genome sequencing (WGS) provides sufficient resolution to distinguish between strains. Expanding on the previously described BA lineages A, B and C (22), in 2020 Bruce et al. provided a new and more detailed classification framework of BA. Based on WGS data, it comprises six primary clusters separated into 18 nested clades (23), offering much higher resolution. WGS studies have also allowed characterisation of outbreak strains in greater detail than previously possible (24, 25) and reassessment of country-level population structures (26, 27). WGS-informed insights into BA diversity at local geographical scales are still limited but hold great potential to determine transmission patterns and drivers of evolution (28, 29). Previously, we were able to identify 22 BA genotypes within the Ngorongoro Conservation Area (NCA) of northern Tanzania (30), an area of ∼ 8,300 km^2^ that is highly endemic for anthrax (6, 31). These genotypes all clustered within lineage 3.2 as described by Bruce et al. (2020) (23) that falls within the Ancient A sub-lineage as described by Van Ert et al. (2007) (22). Our prior study demonstrated that even at a local geographical scale, BA types can be distinguished based on a small number of single nucleotide polymorphisms (SNPs). This was corroborated by reports of localised BA diversity from multiple anthrax-endemic regions, such as Vietnam (9), the ‘anthrax belt’ in Australia (4) and the Edwards Plateau in Texas in the United States (32). However, the extent to which the level of SNP diversity typically observed in endemic areas allows for inference of epidemiological features, such as epidemiological links or host range restrictions, remains understudied.

For bacterial WGS, isolates are typically cultured in order to obtain DNA of sufficient quality and quantity. However, culture of BA can be challenging due to high biosafety requirements, and appropriate laboratory facilities may not be available in endemic settings. Moreover, it may be difficult to successfully culture the bacterium from samples containing mixed bacterial species, and from older archived samples. To overcome these challenges, we previously developed a targeted sequence capture (TC) method to enable genomic data to be generated directly from samples without the need for culture (33), thereby maximising the use of a range of sample types. Whether TC methods could realistically replace traditional culture-based approaches for genomic epidemiology has yet to be explored.

Our understanding of the epidemiology of neglected zoonoses such as anthrax is often limited by a lack of coordinated data collection across the human-animal-environment continuum (34, 35), making it difficult to assess its inter-species transmission dynamics (36), e.g., as evidenced by strain-sharing. Consequently, key questions regarding the transmission of BA at the One Health interface remain, i.e. in areas where humans, livestock and wildlife share a common environment. For instance, our understanding of biological (e.g., host species, bacterial genotypes) and non-biological (e.g., landscape, climate) factors that impact BA dynamics, evolution and population structure is still limited. While we know that BA can infect many host species (37), certain species may be disproportionately affected during outbreaks despite similar apparent exposure opportunities (38–40). This phenomenon could be linked to genetic differences between strains or to host behavioural traits, as discussed in previous studies (7). Experimental studies have shown that mutations in BA chromosomal and plasmid genes can alter virulence for different host species as well as the bacterium’s capacity to withstand nutritional stress during infection (41–43). Both of these factors could influence infection success depending on the host species and the characteristics of its immune system. Host predilection can be investigated by searching for existence of host-associated genotypes that disproportionately affect certain host species (44). This in turn could help to assess anthrax risks, such as if certain strains are more likely to infect humans or specific livestock species, representing distinct reservoirs.

This study focuses on the NCA as an example of an area known for frequent cases and outbreaks of anthrax in people, livestock, and wildlife (6, 10). The NCA – part of the Arusha Region, bordering the Serengeti National Park – is a protected zone where a variety of wildlife species and human pastoral communities have shared the same environment for centuries (45). As the southern African region is suspected to be the evolutionary origin of anthrax (22), the disease has likely been endemic in Tanzania for hundreds, if not thousands of years (22, 46, 47).

We hypothesised that multiple anthropogenic and ecological factors could influence the population structure of BA. In the NCA, pastoral livestock holders guide their animals along extensive daily routes for grazing and watering. In so doing, animals are regularly exposed to areas that local communities understand, and microbiologic investigations confirm, pose an increased risk of anthrax but where resource limitations make it challenging to avoid (46). These identified areas of perceived high risk for infection, sometimes referred to as anthrax hotspots, are often associated with proximity to watering sites and human settlements. Livestock are most likely to die and deposit spores in the areas that they most commonly frequent. Although grazing routes are extensive, daily movements usually remain within ∼5 km of livestock-keeping households.

Environmental factors such as fertile soil, dense vegetation, and bodies of water attract hosts and give reason to establish settlements or for herdspersons to lead their animals into areas with such features (7, 14, 48). Additionally, spore formation success in soil is higher under humid compared with dry conditions (19, 49), which might further increase anthrax events during rainy seasons or in areas with denser vegetation and levels of organic matter in the soil (50). Currently there is a paucity of data on whether areas of higher infection risk, or factors such as location of settlements or optimal grazing areas, may lead to the accumulation of genetically related strains within these areas.

While the seasonality of anthrax disease is well-documented globally, the relationship between weather conditions and disease occurrence varies by geographic region (39, 51, 52). The NCA experiences two wet and two dry seasons (53), with the rainy seasons boosting vegetation growth. The temporal limitation in how long some areas may be attractive to hosts during the year could limit the access to spore deposits in those areas. Conversely, flooding during the rainy seasons can unearth and spread spores along riverbanks (54), expanding and reshaping the area in which specific strains circulate over time. In the NCA, communities report seasonal peaks in anthrax cases, particularly during the longer dry season (55). To date, the interplay between seasonality, environmental risk, and genetic relatedness of BA strains in the NCA has not been explored.

Our study had two major objectives. First, we aimed to characterise the genomic diversity of BA in the NCA at the human-livestock-wildlife interface, as an example of a highly endemic area, by assessing the frequency and spatial-temporal distribution of SNP profiles. As part of this objective, we sought to assess whether our culture-free TC-derived sequences offer qualitatively similar conclusions regarding BA diversity as isolate-derived sequences, including through the examination of sequences derived from the same carcass or sample using both methods. Second, we aimed to test whether BA exhibited genetic population structure and to look for evidence of trait-specific BA lineages. Specifically, we tested for phylogenetic clustering by host type (livestock/wildlife), host species, season, and detection within high-risk zones for anthrax and within epidemiologically linked groups. For both objectives, we examined whether the extent of diversity is structured across nested physical scales: i) between cases, ii) within a single case and iii) within the same sample (e.g. tissue) within a case.

## Methods

### Ethics Statement

This study received ethical approval from the Kilimanjaro Christian Medical University College Ethics Review Committee (certificate No. 2050); the National Institute for Medical Research, Tanzania (NIMR/HQ/R.8a/Vol. IX/2660); Tanzanian Commission for Science and Technology (2016-95-NA-2016-45); and the College of Medical Veterinary and Life Sciences ethics committee at the University of Glasgow (200150152). Written consent was obtained from all human participants. Human samples from Moshi were collected under KCMU Co EC certificate #295, NIMR/HQ/R.8a/Vol. IX/1000.

### Sample collection

Samples and associated metadata were collected in the NCA from 2016 through 2021. Our field team were informed about suspect anthrax cases (i.e., people with symptoms and signs of anthrax, and carcasses of animals that died suddenly without prior signs of illness or predation) by NCA community members, as described in more detail elsewhere (6). For livestock and wildlife carcasses, the following samples were collected depending on their availability: blood, tissue, swabs from open orifices, and soil from underneath and insects in close proximity to the carcass, as previously described (30, 56). Samples were collected as soon as possible following the death of the animal, typically within a few days, although occasionally up to a few weeks. Where possible, metadata were documented on host species, Global Positioning System (GPS) coordinates of the sampling location, and date of sample collection. For humans with suspected cutaneous anthrax, swabs were taken from the liquid below the eschar by trained health professionals. Samples from a human with neurological signs (among other signs) confirmed to have died of anthrax from near Moshi, Kilimanjaro Region, were available through the Kilimanjaro Christian Medical Centre; these were provided as DNA extracts (n = 2) from a brain tissue sample. Additional DNA extracts (n = 5) from animal (4) and human (1) blood samples that had tested positive for BA by qPCR– collected in the Arusha Region of Tanzania from areas outside the NCA from 2015 through 2023 – were made available through surveillance efforts led by the Tanzania Veterinary Laboratory Agency (TVLA) supported by the collaborative biosafety project between TVLA and the Finnish Centre for Military Medicine (SOTLK).

### DNA extraction and sequencing preparation for bacterial isolate culture

Fifty-one samples were cultured on sheep blood agar at the Kilimanjaro Clinical Research Institute (KCRI), Moshi, Tanzania. DNA was extracted from colonies grown overnight at 37 °C, using either the QIAamp UCP Pathogen Mini kit or the DNeasy Blood & Tissue kit (Qiagen). The final DNA eluate was filtered through a 0.2 μm column, centrifuging at low speed (1000 rpm) for 4 minutes to eliminate any residual spores. These resulting DNA extracts from isolates – confirmed to be positive for BA by qPCR as previously described (56)– were subjected to library preparation and Illumina sequencing at MicrobesNG (Birmingham, UK).

### Target-enrichment sequencing

One hundred eleven TC sequences were generated for this study following protocols previously described in (33). Briefly, library preparation was carried out using the NEBNext Ultra II FS DNA Library Prep Kit for Illumina, according to New England BioLabs manual Version 3.0_09/22 (57). RNA baits were used to selectively capture the core genome of BA as described (33). Briefly, baits of approximately 80 base pairs (bp) in length were designed to achieve ∼50% overlap across the BA core chromosomal genome, ensuring that at least two baits covered any given position, and excluding plasmids (PXO1 and PXO2). This core genome was determined using 52 BA genomes publicly available at the time, which included the three major clades (22) and covered 89% of the Ames Ancestor sequence NC_007530.2 (58). Hybridisation for BA target enrichment was carried out according to the myBaits Hybridization Capture for Targeted NGS manual v4.01 (majority) or v5.02 (samples provided by TVLA) (59), using the Standard protocol.

All samples from the NCA processed with TC were sequenced on an Illumina NextSeq 500 platform, generating read lengths of 2x75 or 2x100 bp. Among the archived blood samples from outside the NCA available through TVLA, seven DNA extracts were initially selected for TC, based on having a qPCR cycle threshold (Ct) value below 35 for at least one BA-specific target, suggesting higher concentrations of BA-specific DNA. For these samples, sequencing preparation of hybridisation captured-libraries was carried out according to the Denature and Dilute Libraries Guide manual for the NextSeq System, Version of December 2018 (60) following Standard Normalization protocol and Best Practice guidelines for Standard and Bead-Based Normalization in Nextera XT DNA Library Preparation Kits (61) and sequenced on an Illumina NextSeq 550 platform, generating read lengths of 2x150 bp.

### Read mapping and SNP calling

Raw reads from a total of 288 samples were processed in a custom pipeline for read mapping and SNP calling. In addition to the sequences generated in this study, 72 isolate-derived sequences from (30) and 52 TC-derived sequences from (33) meeting the minimum coverage described below were included. The original pipeline and a Nextflow version are available on GitHub (https://github.com/matejmedvecky/anthraxdiversityscripts; https://github.com/tlforde/OHRBID). In brief, raw reads were trimmed using Trimmomatic (v. 039) (62), and mapped against the Ames ancestor reference sequence (NC_007530.2) (58), using Bowtie v.2 to ensure mapping of full rather than partial reads and setting an insert size of 800 bp (63). Varscan (64) was run with a coverage threshold of 8 (computing the variance at all positions with at least 8 reads) and a minimum variant frequency of 80% to call variant sites. This value was selected because higher values for homozygosity at a coverage threshold of 8 reads would have led to an unreasonable loss of data. Read mapping metrics across all samples for SNP sites only are created within a second step, using the bam-readcount tool. From initially called SNP sites, only base alleles with a minimum of 6X read coverage and a minimum homozygosity of 80% were called; this threshold was selected with the goal of calling fixed alleles and minimizing artefacts introduced through sequencing error. Moreover, all sites in which minimum read coverage and/or alternate allele frequency thresholds were not met by ≥20 samples were removed. Remaining sites below our coverage threshold were denoted as gaps (‘-’) and sites below frequency threshold were denoted as ambiguous (‘N’) in the resulting multiple sequence alignment file that was used for the subsequent phylogenomic analysis.

Out of all collected samples (n = 288), a total of 75 sequences were eventually excluded due to failing to meet the coverage criteria (mapped reads had to cover > 79% of the Ames Ancestor reference genome at ≥ 8X average depth of coverage) or sample contamination identified by manual inspection of BAM files with suspicious frequencies of alternate alleles via Tablet v1.21 (65). The final alignment of concatenated SNPs included 213 sequences (122 derived from WGS of cultured isolates, and 91 from TC; **Error! Reference source not found.**), with a total length of 1196 bases.

### Phylogenomic analysis

Model selection and maximum likelihood (ML) tree estimation were performed in IQtree (66) using slow bootstrapping with 1,000 replicates, which suggested a Kimura-3-parameter model with equal base frequencies. As the alignment included only polymorphic sites, invariant positions were excluded. This leads to ascertainment bias when inferring phylogenetic models, as standard substitution models assume the presence of both variable and invariant sites. The IQTree option of ascertainment bias correction was therefore applied to adjust likelihood calculations accordingly.

Tree samples for clustering assessment were generated with MrBayes Version 3.2.7a (67) and applying the Kimura-3-parameter substitution model. Analyses were run using an unconstrained branch length prior at a chain length of 10,000,000 iterations, sampling every 10,000 states and removing 10% burn-in. The full phylogeny was generated using the dataset of 213 high-quality sequences and the Ames ancestor reference as an outgroup, as it represents a different sub-lineage from Ancient A (58).

To test for phylogenetic clustering (described below), the dataset was de-duplicated by case, such that if multiple identical sequences per case were sequenced, the sample with the highest coverage was selected. This resulted in an alignment of n=150 genomes.

### Identifying unique SNP profiles and SNP distances

A set of unique ‘SNP profiles’ present within the data was defined, wherein each SNP profile differs from all others by at least one SNP (as opposed to any differences arising from gaps or ambiguous base signal). After collapsing unambiguous duplicates using a custom R-script, a secondary identity check using pairwise genetic distances using the dist.dna function (R package ape) further identified minor nucleotide differences among sequences that passed the first filter, resulting in a final refined set of unambiguously distinct SNP profiles (68–70). Specifically, within the dist.dna function, we used the Hamming distance which calculates to the absolute number of differing sites (for simplicity henceforth referred to as SNP distance). Sites with ambiguous base signal and gaps were excluded in a pairwise manner throughout rather than across the full dataset to maximise the information retained. Beyond phylogenetic approaches, we further used the mean Hamming distance (SNP distance) of groups of sequences, considering the similarity between sequences as another aspect of diversity and relatedness. SNP distances among different groups of sequences were defined and compared, as described below.

### Testing the effect of sequence retrieval method on detected diversity

We assessed whether sequences retrieved via TC and isolate culture showed a similar level of diversity using three approaches. Our primary aim was to test whether the metagenomic TC approach might introduce some level of spurious (artefactual) diversity. First, we conducted a rarefaction analysis using the iNEXT package in R (71) comparing the 122 sequences (55 cases) obtained via isolate culture with all 91 sequences (84 cases) obtained via TC. Second, a permutation test was performed to investigate whether there was a significant difference between the mean SNP distance among sequences retrieved via isolate or TC, while accounting for sample size differences in the number of cases. To this end, the same number of cases (n = 55) from which all isolate sequences had been collected from was sub-sampled from those 78 TC cases (only those that had been found within the NCA). This was repeated to produce 100 TC-data subsets. We tested for a difference in mean SNP distance among each of the 100 subsets and the full set of isolate sequences. An initial α threshold of 0.05 was Bonferroni corrected to 0.0005 to account for repeated testing. We also assessed instances in which multiple genomes had been sequenced from the same sample material using both methods to verify whether sequences derived by TC and isolate culture would show the same SNP profile. Third, we assessed to what extent a higher number of gaps and ambiguous sites among TC-derived sequences, which are ignored during pairwise sequence comparison, would lead to an under-estimation of diversity. We used Bali-phy (v.4.0-beta.16 (72)) to infer ancestral character states of a fixed alignment, with a general time reversible substitution model. Two independent runs of 50,000 states were combined, and a consensus tree and alignment were determined using the bp-analyze script (https://www.bali-phy.org/).

### Genomic distances and diversity on multiple physical scales

Because both diversity and population structure may emerge at different physical scales, we examined SNP distances among cases, within individual cases and within sample material to assess whether SNP distances and diversity decrease towards smaller scales. For the calculation of within case and between-case genomic distance, sequences were grouped by case and mean distances for each case and between case groups were determined in a pairwise manner. For the assessment of within-sampling material diversity, only sequences derived from tissue and blood could be definitively considered the same material. Other sample types from a carcass site were routinely collected in aliquots that would mean that isolates or DNA extracts could effectively come from different body sites (e.g. for multiple swabs taken), and these samples were therefore not included in this analysis.

### Assessing geographic and epidemiological traits with respect to *Bacillus anthracis* population structure

Cases were considered to be putatively epidemiologically linked (epidemiologically linked groups, ELGs) if they occurred within close spatial-temporal proximity (occurring within 10 days and within 10 km of each-other) (see Microreact project: https://microreact.org/project/ivGWEG2R5UUM3vdQ5kpTck-ba-ncahilbig-medvecky-forde). These values were chosen based on the common incubation time for anthrax of 3-5 days (19) and the average daily distance of 4.26 km travelled by pastoralists with their herds in the NCA (46). We documented the distribution of these ELGs and their size in terms of the number of SNP profiles, sequences and cases within them. Zones of high and low risk for anthrax infection in livestock within the NCA were previously determined by participatory mapping studies (46) using a composite model that combined landscape and climatic factors; see supplementary Figure S1 for a schematic representation of the high-risk area, which covers roughly 700 km^2^.

We formally compared genomic (SNP) distances for sequences grouped by different covariates. Specifically, we compared mean SNP distances between groups of sequences sampled from epidemiologically linked cases to those not linked, and between sequences from cases occurring in areas of high versus low risk of infection. To this end, permutation tests were conducted for significance testing of comparisons (sequences ELGs vs. not epidemiologically linked and high vs. low infection risk areas) by randomly reshuffling value assignments while preserving category proportions. The difference between means was calculated 10,000 times.

### Assessing population structure through phylogenetic clustering by trait

We wanted to understand whether certain non-biological and biological factors place limitations or pressure on the circulation of strains, either geographically (e.g., whether strains concentrate in certain areas, potentially further affected by climate) or in terms of host predilection (e.g., whether some strains are associated with certain types (i.e. wildlife versus livestock) or species of animals. We tested for this by determining whether certain traits were associated with stronger phylogenetic clustering. We assessed this for i) host type (livestock/wildlife), ii) host species, iii) season of anthrax occurrence, iv) collection sites located within vs. outside a high-risk zone and v) putative ELGs.

Variables for season were informed by prior research conducted in the NCA (55): four seasons differing by average temperature and levels of precipitation had been described in focus group discussions involving NCA community members. Participants identified historic peaks in anthrax occurrence between January and March (short dry season) and June to October (long dry season), in line with other studies from the region (10, 53). However, cases are reported throughout the year, including during April and May (long rains) and November and December (short rains). In our analyses, data for each season were combined across sampling years.

To assess phylogenetic clustering of these five categorical traits, a UniFrac-based method (73, 74) that could account for clustering on the trait state level and include phylogenetic (branch length) information was used, as implemented within the SeqTree toolkit (https://github.com/will-harvey/toolkit_seqTree). This function computes a tree-aware clustering score per trait level, with higher values indicating tighter phylogenetic grouping. For each trait, 100 annotated phylogenetic topologies derived from MrBayes posterior samples (see above, section ‘Phylogenomic analysis’) were analysed. Trait labels were shuffled 1,000 times across tip nodes per tree to generate null distributions of UniFrac scores. Observed UniFrac values were compared to null distributions to calculate empirical right-tailed p-values, followed by false discovery rate correction using the Benjamini–Hochberg method. All permutation analyses were performed in R. Results were then summarised across the 100 trees using a custom Python script.

### Data Availability

All sequence data generated as part of this study are available under BioProject accession numbers PRJNA918617 (isolates) and PRJNA1298849 (targeted capture). Individual BioSample accessions are also included in the Microreact project spreadsheet. The following metadata are provided in the Microreact project: https://microreact.org/project/ivGWEG2R5UUM3vdQ5kpTck-ba-ncahilbig-medvecky-forde: cases, species, type of host, sampling location and year, classification of sampling location as high or low risk for anthrax infection, as well as classification of sampling time during the year into one of four typical seasons for the occurrence of anthrax cases, sequence retrieval method, epidemiologically linked groups, SNP profiles as found within this study and in comparison to the genotypes defined in the predecessor study, sample material, as well as the study in which any sequences were originally published. A read-me file about the data provided within the Microreact project is available with the Supplementary Material 2.

## Results

### Genomic dataset

The final dataset of 213 sequences contained 206 sequences from within the NCA, five TC-derived sequences were from the Arusha region outside of the NCA, and two TC-derived sequences were from the Moshi-based decedent. The sequences were derived from 121 anthrax cases from eight different species. Forty-four cases were represented by more than one sequence (range: 2-6, median 3).

### Genomic diversity of *Bacillus anthracis* in an endemic area

Phylogenetic reconstruction of the 213-sequence dataset from the NCA and neighbouring areas (Figure 1) revealed that all sequences sampled within the NCA fell into a main “NCA-clade”, corresponding to cluster 3.2 as described by Bruce et al. (2020) (23). In contrast, all seven sequences sampled outside the NCA but from the wider Arusha Region and Moshi, fell into two “non-NCA-clades”, both of which belonged to the 3.1 cluster. One sequence showed an ambiguous pattern of allele states at the sites of the cluster-defining SNPs (Supplementary Material 1, section 1; Table S1).

**Figure 1:**
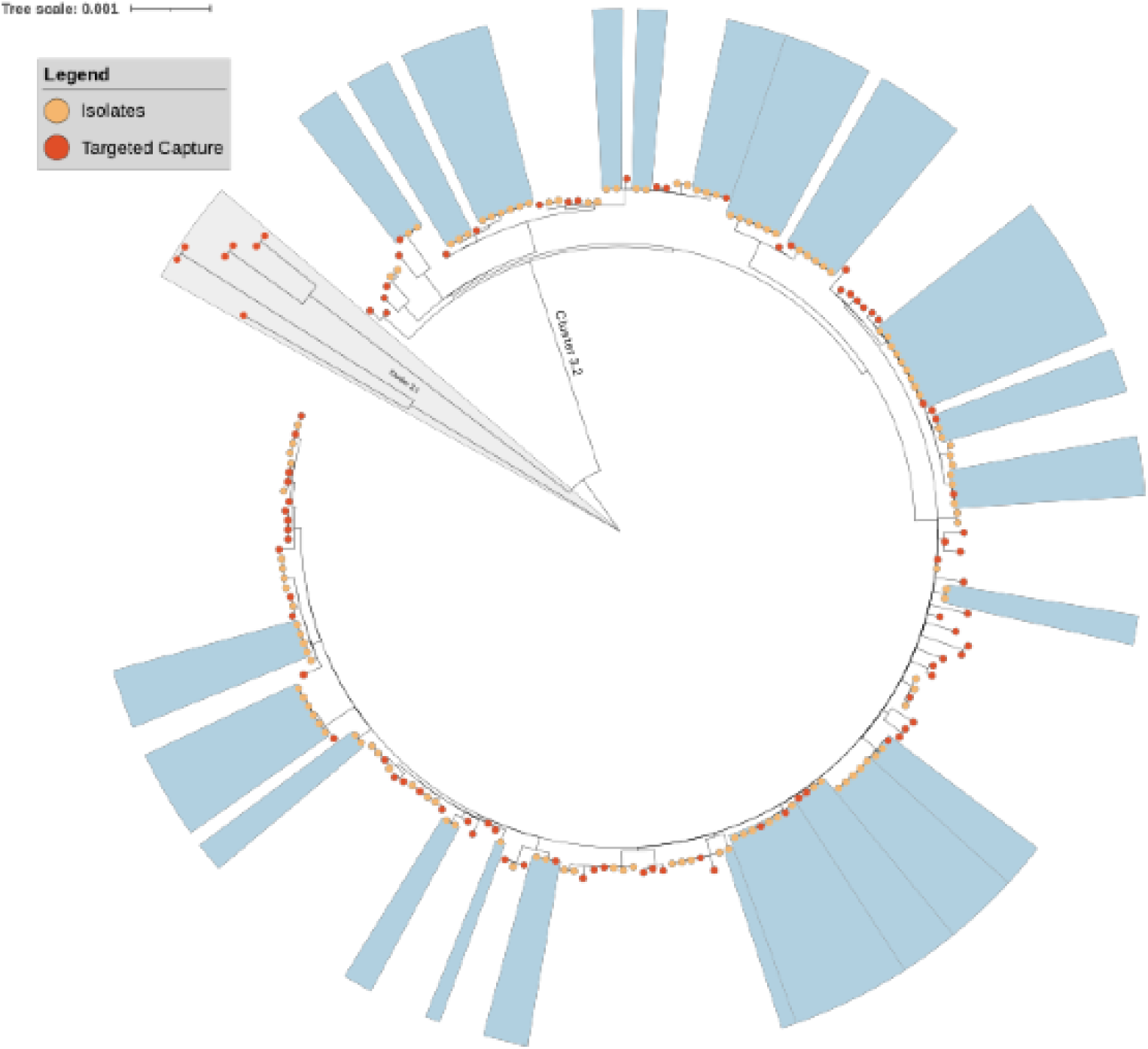
Maximum likelihood phylogeny of Bacillus anthracis (BA) genomes from the Ngorongoro Conservation Area (NCA) and broader Arusha and Kilimanjaro regions analysed in this study (n = 213). The tree was estimated using IQTree. BA genomes were collected between 2015 – 2023. The Ames Ancestor genome is used as an outgroup for rooting (not shown). Blue panels represent sequences belonging to the 22 single nucleotide polymorphism (SNP) profiles previously described by Forde et al. (2022) (30). Coloured circles show sequences derived from isolates (peach) or targeted capture (red); those not falling within the blue panels represent newly-identified SNP profiles. All NCA sequences fell within cluster 3.2 defined by Bruce et al. (2020)(23), whereas sequences from outside the NCA clustered distinctly into separate clades, corresponding to the 3.1 cluster. An annotated version of this figure is provided as supplementary Figure S2.

Among 213 BA sequences, we observed 104 SNP profiles that differed by at least one SNP. Of these, 22 had been previously described using isolate-derived sequences (30), while 82 were newly detected (Figure 1; Figure S2). In the current expanded dataset, we observed 14 sequences belonging to those 22 previously defined SNP profiles, nine of these sequences were retrieved using TC and five from cultured isolates. Among the 104 SNP profiles detected, 47 were observed in at least two separate samples and 24 SNP profiles in at least two different cases. Of these 24 SNP profiles shared between cases, 18 were found in cases >10 km and at least two weeks apart from one-another (i.e. not epidemiologically linked). There were ten SNP profiles that were detected in cases that occurred >6 months apart. Of these, six were detected in case pairs, while the remaining four were sampled from each 4-6 cases that occurred over time intervals of 7 months to 2 years. Notably, the same SNP profile was identified in samples collected from two cases in the Arusha Region outside the NCA that occurred over three years apart. The SNP profiles of highest prevalence among cases were found in five and six separate carcasses, respectively. Only two of these cases were putatively epidemiologically linked.

### Impact of targeted capture versus culturing on the detected diversity

Of the 104 SNP profiles, 57 were represented from TC-derived sequences alone, 28 amongst isolate-derived sequences alone, and 19 were represented by both methods. A rarefaction analysis showed that sampling of isolate sequences likely led to good coverage of an extrapolated diversity (Figure 2A), whereas this was not the case for sampling results of TC sequences (Figure 2B). This finding was observed regardless of whether TC derived sequences were limited to those within the NCA clade or included all sequences. We found 47 (39%) unique SNP profiles among 122 isolate-derived sequences and 76 (84%) unique SNP profiles among 91 TC-derived sequences.

**Figure 2:**
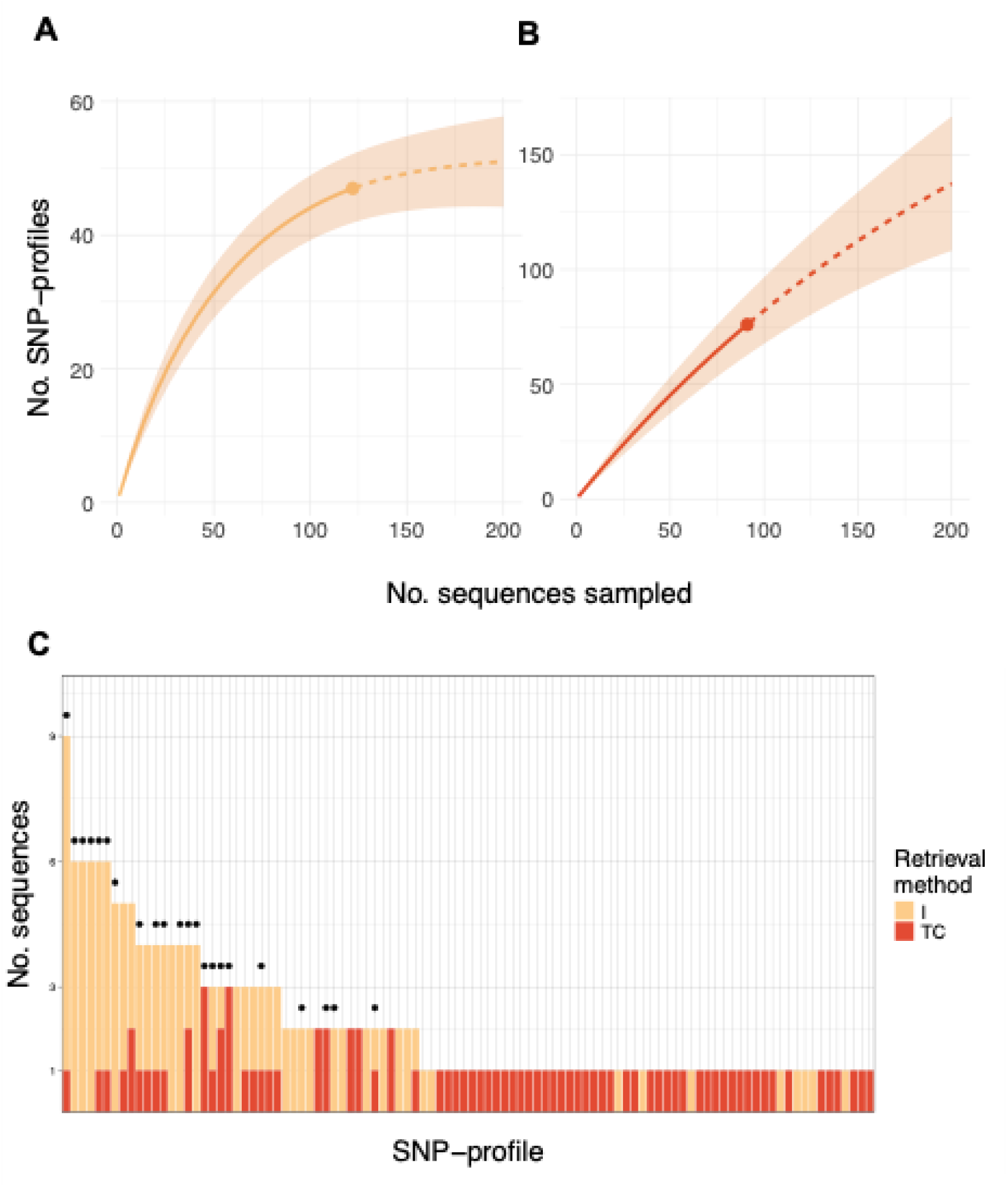
Rarefaction analyses and distribution of abundance for Bacillus anthracis single nucleotide polymorphism (SNP)-profile diversity comparing sequences derived from isolate culture and targeted capture (TC). Retrieval method is indicated by colour. Rarefaction analyses estimating how successfully the existing diversity has been sampled, comparing between sequences derived from isolates and TC (A and B, respectively). While the curve for isolate-derived sequences levels off, indicating that all or most SNP profiles have successfully been sampled among the 122 sequences, the curve for TC-derived sequences (n=91) indicates that many more samples would be required to sufficiently capture the SNP profile diversity. Panel C shows all 104 unique SNP profiles and the frequency of their detection, lacking an evident pattern in their distribution. SNP profiles collected from more than one case are marked by a black dot above the bar.

SNP profiles were represented by as many as nine sequences and there was no distinct pattern in their abundance (Figure 2C). Of 44 SNP profiles represented by more than one sequence, in 22 instances these profiles were sampled from the same case, while the other 22 were sampled from multiple cases (Figure 2C). It should be noted that the number of SNP profiles represented by more than one sequence and the number of cases for which multiple genomes were sequenced are both 44 by coincidence.

Of 44 cases from which multiple sequences were generated, 18 comprised sequences derived from both methods. Among the 18, in 15 at least one pair of a TC and an isolate sequence were assigned the same SNP profile. No case contained more than one SNP profile represented by both methods.

SNP distances among all isolate-derived sequences (mean 19.2, median 18 and interquartile range [IQR] = 20) were compared against those observed among sequences from 100 TC-derived sequence subsets from the NCA clade using a permutation test. The number of TC derived sequences per data subset representing 55 cases ranged from 57 – 62 among the 100 subsets. Results were very mixed, with a range of observed differences between -3.4 and 0.6 for the means of SNP distance values of TC to isolate-derived sequences, and a median of -1.1. This indicates a trend towards slightly larger SNP distances among sequences derived from isolate culture, however these distances ranged between 1-2 SNPs. The permutation tests yielded significant p-values (after Bonferroni correction of the α threshold from 0.05 to 0.0005 for 100 tests) for 51 of 100 data subsets and all of these were associated with negative observed differences, indicating larger SNP distances among isolate derived sequences; however, 41 of 100 p-values were associated with negative observed difference in means but were non-significant (see supplementary Figure S3).

For four cases, we were able to obtain both isolate and TC derived sequences from the same tissue sample (Table 2). In three of these instances, isolate- and TC-derived sequences from the same case belonged to the same SNP profile; in the two instances where the SNP profile of a third sequences differed from the other two, one was a TC and one an isolate-derived SNP profile, each differing by one SNP.

**Table 1:** Anthrax cases and associated Bacillus anthracis sequences (n=213) from northern Tanzania, structured by sequence retrieval method and host species. Total counts are given for unique cases represented by samples processed with either retrieval method. Number of sequences are shown in parentheses.

|  | Elephant | Wildebeest | Zebra | Cattle | Donkey | Goat | Sheep | Human | Not recorded | Total |
| --- | --- | --- | --- | --- | --- | --- | --- | --- | --- | --- |
| **Total cases** | 1 | 8 | 4 | 19 | 1 | 14 | 62 | 6 | 6 | 121 |
| **Isolate** | 0 | 4 (11) | 3 (7) | 8 (22) | 1 (2) | 6 (16) | 27 (57) | 1 (1) | 5 (6) | 55 (122) |
| **Targeted Capture** | 1 (1) | 7 (9) | 4 (6) | 13 (13) | 1 (1) | 11 (11) | 41 (43) | 5 (6) | 1 (1) | 84 (91) |

**Table 2:** Assessment of single nucleotide polymorphism (SNP)-profiles between sequences from the same tissue that were derived by either isolate or targeted capture (TC) method. . In three of four cases, SNP profiles match in sequences derived from both methods.

| Case | Method of obtaining sequences (same tissue) | No. sequences | Sample material | Sequences matching |
| --- | --- | --- | --- | --- |
| 1 | 1 Isolate, 1 TC | 2 | Blood | Match |
| 2 | 1 Isolate, 1 TC | 2 | Tissue | Match |
| 3 | 2 Isolates, 1 TC | 3 | Tissue | TC derived SNP profile matches one isolate, SNP profile of the second isolate differed by 1 SNP |
| 4 | 2 Isolates, 1 TC | 3 | Tissue | SNP profiles of isolates match, SNP profile of TC differed by 1 SNP |

We observed a higher number of missing and ambiguous SNP sites among TC sequences. However, the difference in median values for both types of sites between the two methods was low compared to the difference in mean and maximum values (Table 3). A visual representation of these distributions can be found in supplementary Figure S4.

**Table 3:** Occurrence of sites with missing or ambiguous allele information among Bacillus anthracis sequences derived from isolates (I, n = 122) and targeted capture (TC, n=91). Range (minimum - maximum) mean and median values for each type of site are given for all sequences derived by either method.

| Method | Gaps |  |  | Ambiguous sites |  |  |
| --- | --- | --- | --- | --- | --- | --- |
|  | Range | Mean | Median | Range | Mean | Median |
| I | 0 - 103 | 7.15 | 1 | 0 - 9 | 0.71 | 0 |
| TC | 1 - 215 | 14.15 | 2 | 0 - 243 | 22.88 | 5 |

Bali-Phy analyses confirmed that diversity could in principle be underestimated due to the presence of gaps and ambiguous characters, but that the extent of this overestimation was very limited. The total number of SNP profiles determined from an alignment in which gaps and ambiguous sites had been inferred only generated a single additional SNP profile, resulting in 105 profiles in comparison to 104 observed.

### Genomic distances and diversity of *Bacillus anthracis* sequences grouped by nested physical scales

Mean pairwise SNP distances were lowest within cases and highest across the full dataset (Table 4). Maximum value results for the full dataset and between cases differ slightly, as pairwise deletion in case-comparison-groups included slightly fewer sites.

**Table 4:**
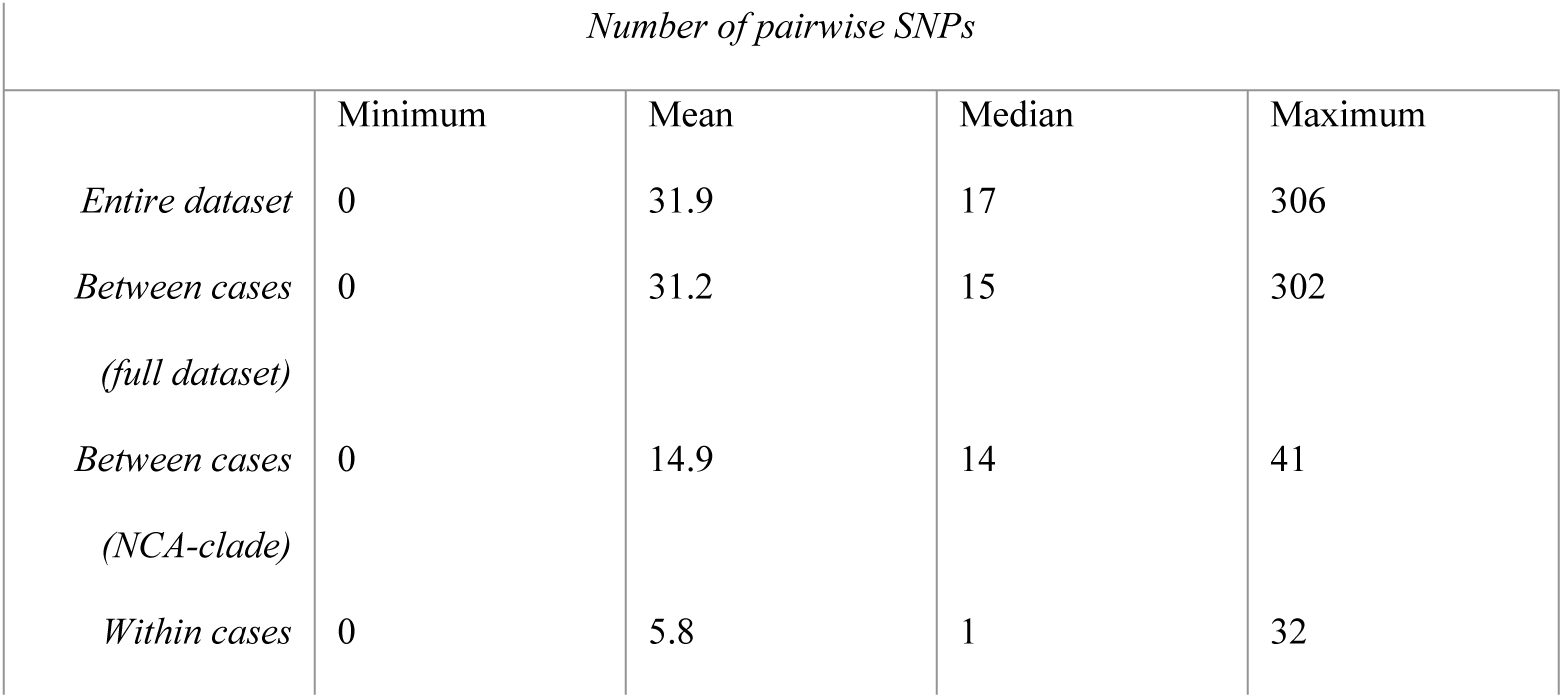
Distribution of mean pairwise single nucleotide polymorphism (SNP) differences between Bacillus anthracis sequences from northern Tanzania (n=213). Values are given for the whole dataset, between all cases, between only those cases collected within the Ngorongoro Conservation Area (NCA) and within cases. Within-case values are given for cases where at least two sequences were available (n_cases_ = 44).

| Number of pairwise SNPs |  |  |  |  |
| --- | --- | --- | --- | --- |
|  | Minimum | Mean | Median | Maximum |
| *Entire dataset* | 0 | 31.9 | 17 | 306 |
| *Between cases*<br>*(full dataset)* | 0 | 31.2 | 15 | 302 |
| *Between cases*<br>*(NCA-clade)* | 0 | 14.9 | 14 | 41 |
| *Within cases* | 0 | 5.8 | 1 | 32 |

Within-case diversity could be assessed for 44 cases from which multiple samples had been collected (2-6 sequences available, median: 3) (Figure 3, Table 4). The majority of cases contained one or two different SNP profiles, many of which were from cases with only two or three sequences in total. No case contained more than three different SNP profiles (Figure 3 B). Considering the full dataset, the largest mean SNP distances between case pairs were driven by the inclusion of the sequences in the 3.1-cluster (non-NCA clade). The largest SNP distances of 259-302 SNPs were found between sequences collected in the Arusha Region, partially from the same year (Figure S5A), while within the NCA clade, between-case SNP distances ranged from 0 to 41 SNPs (Table 4, Supplementary Figure S5B).Within-sample diversity could be assessed for thirteen anthrax cases. Here, 2-3 sequences (median 2) were available from either blood or tissue, respectively, irrespective of sequence retrieval methods. The SNP distances were 0 in eight of thirteen cases. In the remaining five cases, sequences differed by 1 to 28 SNPs.

**Figure 3:**
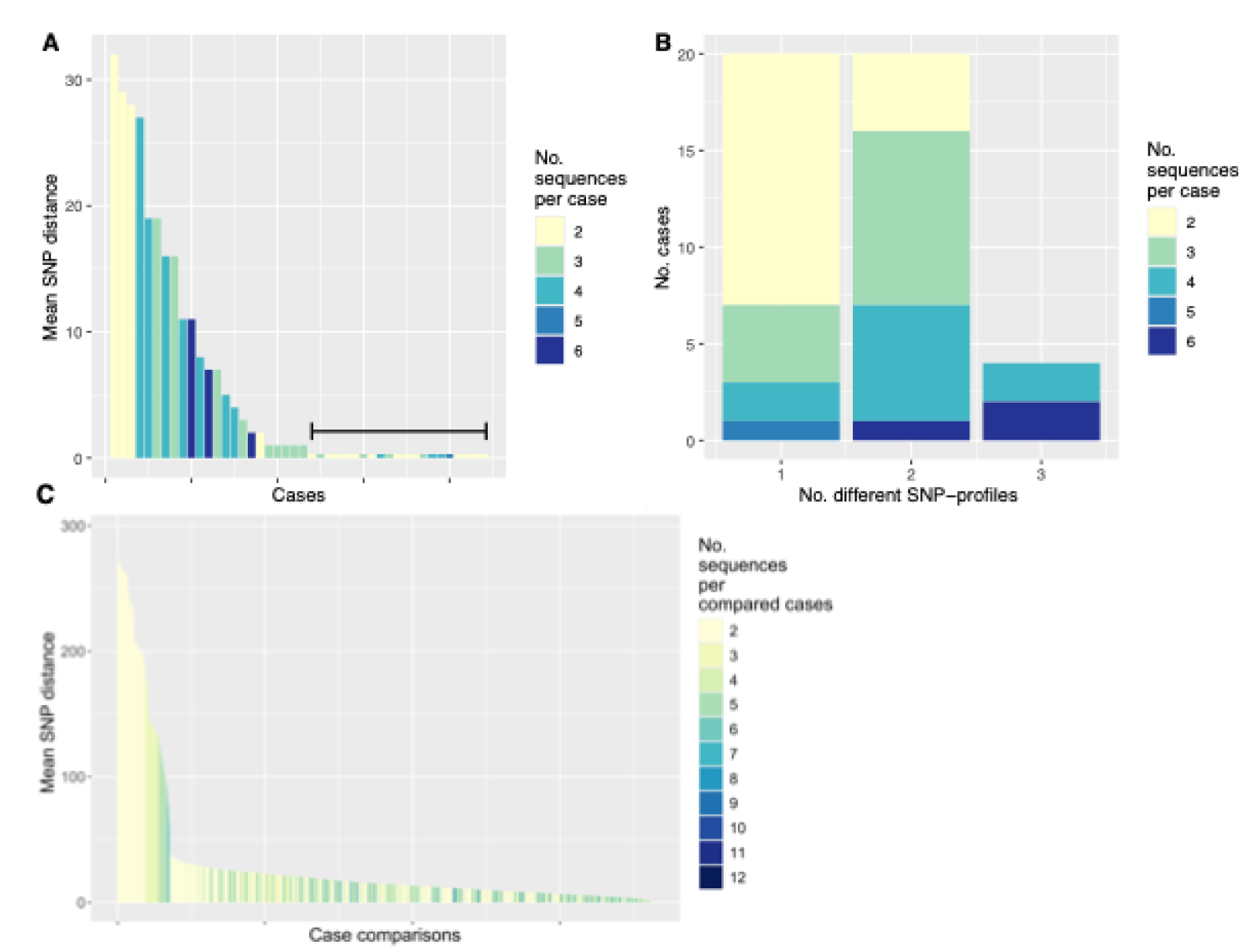
Mean number of single nucleotide polymorphism (SNP) differences between sequences of the full Bacillus anthracis dataset from the Ngorongoro Conservation Area and surrounding areas (n = 213), within and between cases, and counts of differing sequences in cases where multiple sequences were available. Fewer SNP differences separate the sequences from the same cases compared to those retrieved from distinct cases. A) Within cases with more than one sequence (n = 44), mean SNP distance did not correlate with the number of sequences and was highest for cases with sequence pairs. Each bar represents an individual case. For better visualisation, bars for those cases with SNP distance of 0 are raised and demarcated by a black bar. B) Up to three different SNP profiles were identified among sequences from the same case. C) Sequences from the majority of case pairs had SNP distances of < 50, while some differed by over 250 SNPs. Each bar represents an individual case.

### Factors associated with *Bacillus anthracis* population structure

#### Spatio-temporal occurrence of cases and associated SNP distances

There were 22 instances where pairs or groups of cases were sampled within close spatio-temporal proximity and thus considered potentially epidemiologically linked (i.e. within 10 days and 10 km of each other; ELGs outlined in the Methods). Among all 213 samples included in the phylogeny, a total of 121 sequences from 66 cases could be assigned with certainty to one of these 22 ELGs (coloured circles in Figure 4A), which contained 67 different SNP profiles (see Microreact project for details). Of particular interest is ELG-16 (sky blue in Figure 4), which comprised eight anthrax cases involving five wildebeest, two zebras and a sheep, all occurring within 6 days of each-other. Despite their spatio-temporal proximity, these cases harboured a total of 11 different SNP profiles, with two cases (AN18-416 from a zebra and AN18-410 from a wildebeest) sharing a SNP profile (Figure 5; Microreact project).

**Figure 4:**
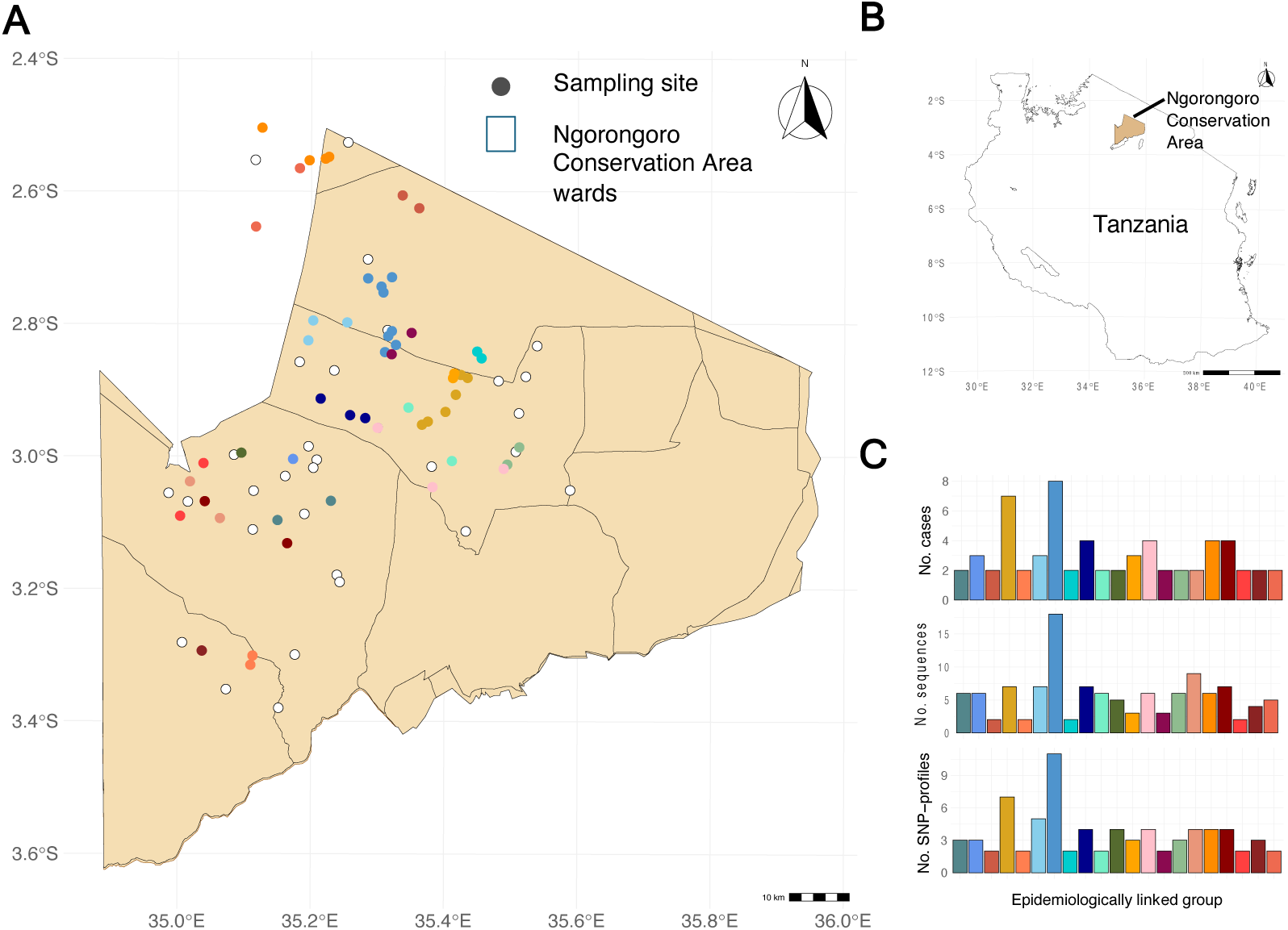
Location of anthrax sampling sites (n = 101) with confirmed Global Positioning System (GPS) data within the Ngorongoro Conservation Area (NCA), northern Tanzania. A total of 177 sequences were obtained from these sites. A) Coloured circles indicate the 64 cases that could be assigned to 21 putative epidemiologically linked groups (ELGs), whereas white circles indicate cases with GPS data but no assignment to an epidemiological group. Administrative wards are shown with a black outline. One ELG, comprising sequences from outside the NCA, is not shown. B) Geographic context of the NCA within Tanzania. C) Barplots showing the number of cases, sequences and single nucleotide polymorphism (SNP) profiles (top, middle and bottom, respectively) within each ELG; colours correspond to those in Panel A.

**Figure 5:**
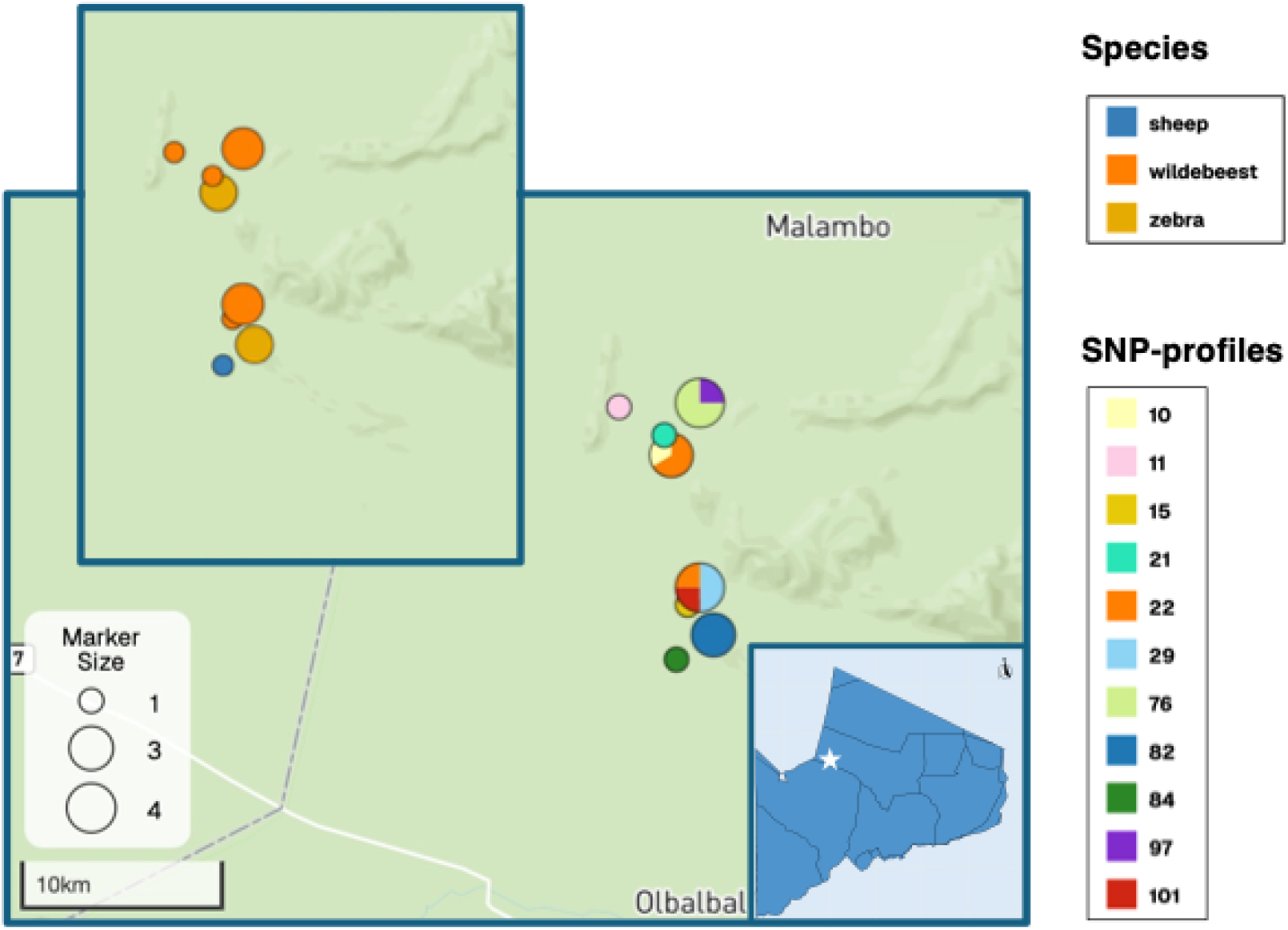
Epidemiologically linked group (ELG)-16, exemplifying the diversity in genotypes that can be observed among anthrax cases despite their occurrence in close spatio-temporal proximity. A total of 18 sequences were obtained from eight cases, representing both livestock and wildlife. Top left: coloured circles represent the species sampled, with size of the circle indicating the number of sequences. Centre: coloured circles indicate the SNP profiles assigned. Bottom right: geographic context, with the white star indicating the location of ELG-16 within Olbalbal Ward, Ngorongoro Conservation Area. The figure is derived from the Microreact project: https://microreact.org/project/ivGWEG2R5UUM3vdQ5kpTck-ba-ncahilbig-medvecky-forde

Pairwise differences were compared among sequences belonging to an ELG (n = 121) versus those without such a link (n = 92) (Table 5). Sequences within ELGs represented 67 different SNP profiles, whereas those without an epidemiological link comprised 53 SNP profiles. On average, the mean SNP distances between sequences not linked to any group were significantly larger (mean 61.3 and median 28 SNPs, IQR 20) than sequences from cases within ELGs (mean 15.8 and median 13 SNPs, IQR 23.8; permutation test, p<0.001). When excluding sequences collected outside the NCA, the difference in means decreased to 6.2 SNPs, yet remained significant (p<0.001, see supplementary Figure S6).

**Table 5:** Distribution of mean pairwise single nucleotide polymorphism (SNP) differences between Bacillus anthracis sequences from northern Tanzania grouped by spatial variables, using the full dataset for sequences sampled in different regions (n= 213). Values are given for sequences from cases of putatively epidemiologically linked groups (ELGs) based on spatio-temporal proximity and for whether the cases occurred within or outside areas perceived by pastoral communities as posing a high risk for anthrax infection.

|  | Number of pairwise SNPs |  |  |  |
| --- | --- | --- | --- | --- |
|  | Minimum | Mean | Median | Maximum |
| **Within ELGs** | 0 | 15.8 | 13 | 301 |
| **Not assigned to ELGs** | 0 | 61.3 | 28 | 306 |
| **Among sequences within the high-risk zone** | 0 | 16 | 13 | 40 |
| **Among sequences outside of high-risk zone** | 0 | 37.7 | 20 | 306 |

In the same way, the diversity of SNP profiles collected outside of the high-risk infection zone was compared to the diversity within. Of 155 sequences found outside the high-risk zone, 89 sequences (57%) were unique. Among 58 sequences collected within the high-risk zone, 37 (64%) belonged to unique SNP profiles.

Mean SNP distances among sequences outside the high-risk zone were larger (mean of 37.7 and median of 20 SNPs (IQR 20) than among sequences within the high-risk zone (mean 16.0, median 13 and IQR 16). The difference in means was significant according to a permutation test (with a p-value of 0). When excluding sequences collected outside the NCA, the difference in means decreased to 3.4 SNPs, yet remained significant (p<0.001, see Supplementary Figure S7).

### Phylogenetic clustering and SNP distances among sequence groups based on host, environmental, or epidemiological traits

We did not observe any apparent phylogenetic clustering based on host species (Figure 6). With the exception of the donkey and elephant, for which sequences were only available for a single case each, all species yielded multiple SNP profiles from throughout the phylogeny. Identical or phylogenetically close sequences were regularly isolated from multiple hosts, including between humans and livestock, and between livestock and wildlife.

**Figure 6:**
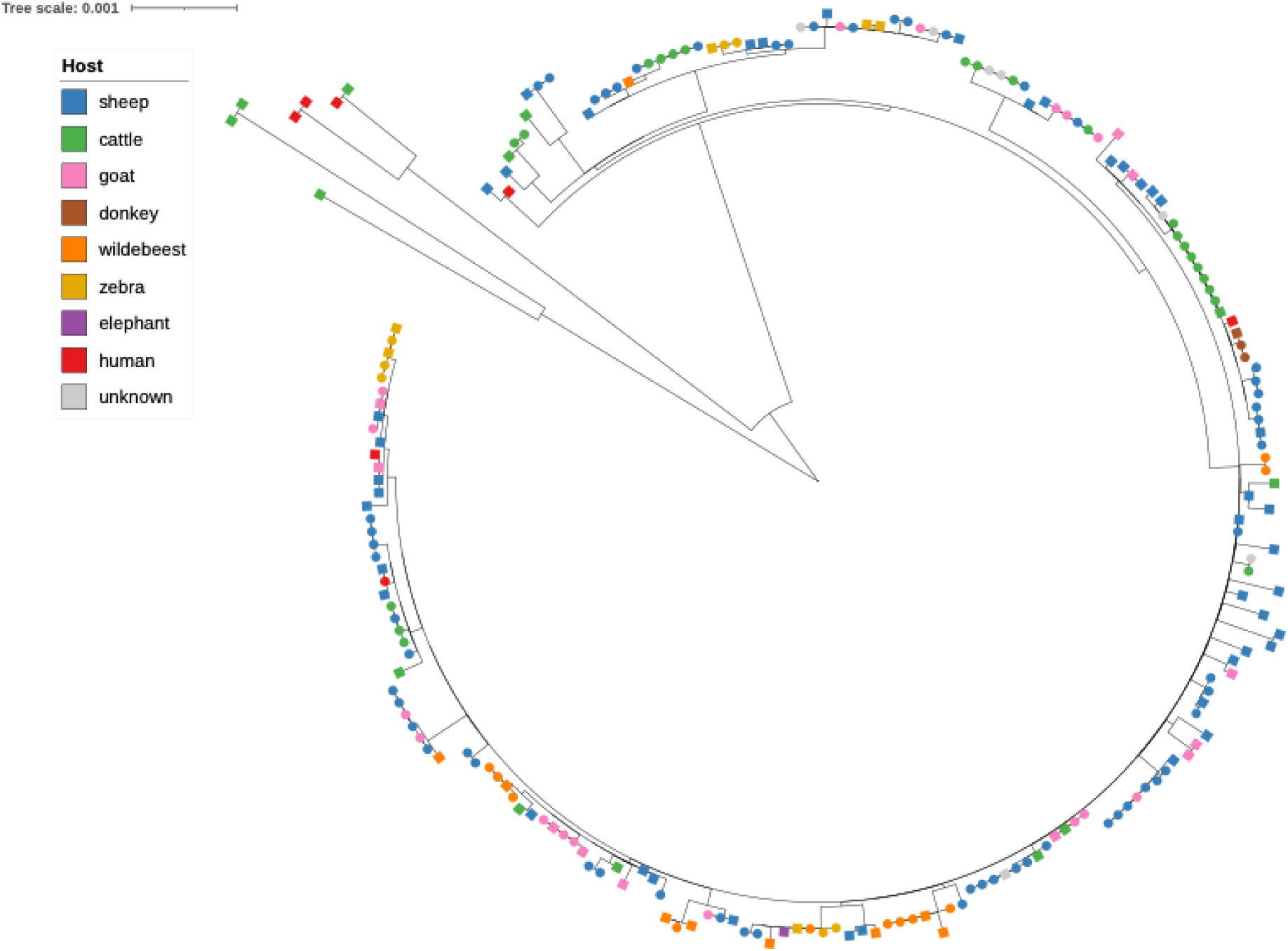
Maximum likelihood phylogeny of Bacillus anthracis genomes from the Ngorongoro Conservation Area and broader Arusha and Kilimanjaro regions analysed in this study (n = 213), highlighting host species of origin. Tree was estimated using IQTree (as in Figure 1). Colours represent different host species, as indicated in the legend. Isolate-derived sequences are shown as circles, whereas those from targeted capture are shown as squares.

Of the five traits formally tested for phylogenetic clustering – namely species, livestock vs. wildlife, season, the ELGs, and sampling location within or outside the high-risk infection zone – two of the traits tested showed a trend towards clustering, yet none of the statistical tests were significant. A single ELG (designated number 15) showed significant clustering (see supplementary Table S2). The trend towards clustering was observed for sequences sampled from sheep and cattle versus other host species (Figure 7A), as well as for sequences from cases sampled in April through May and in November through December (the two wet seasons, long and short, respectively, Figure 7B). No clustering was observed for sequences collected from wildlife versus from livestock. A full list of results can be found in Table S2 in Supplementary Material. The broadly bimodal shape of mean SNP distance distribution in Figure 7 for trait levels cattle, human, Jan-May, Apr-May and Jun-Oct is connected to the groups of sequences from within and outside of the NCA.

**Figure 7:**
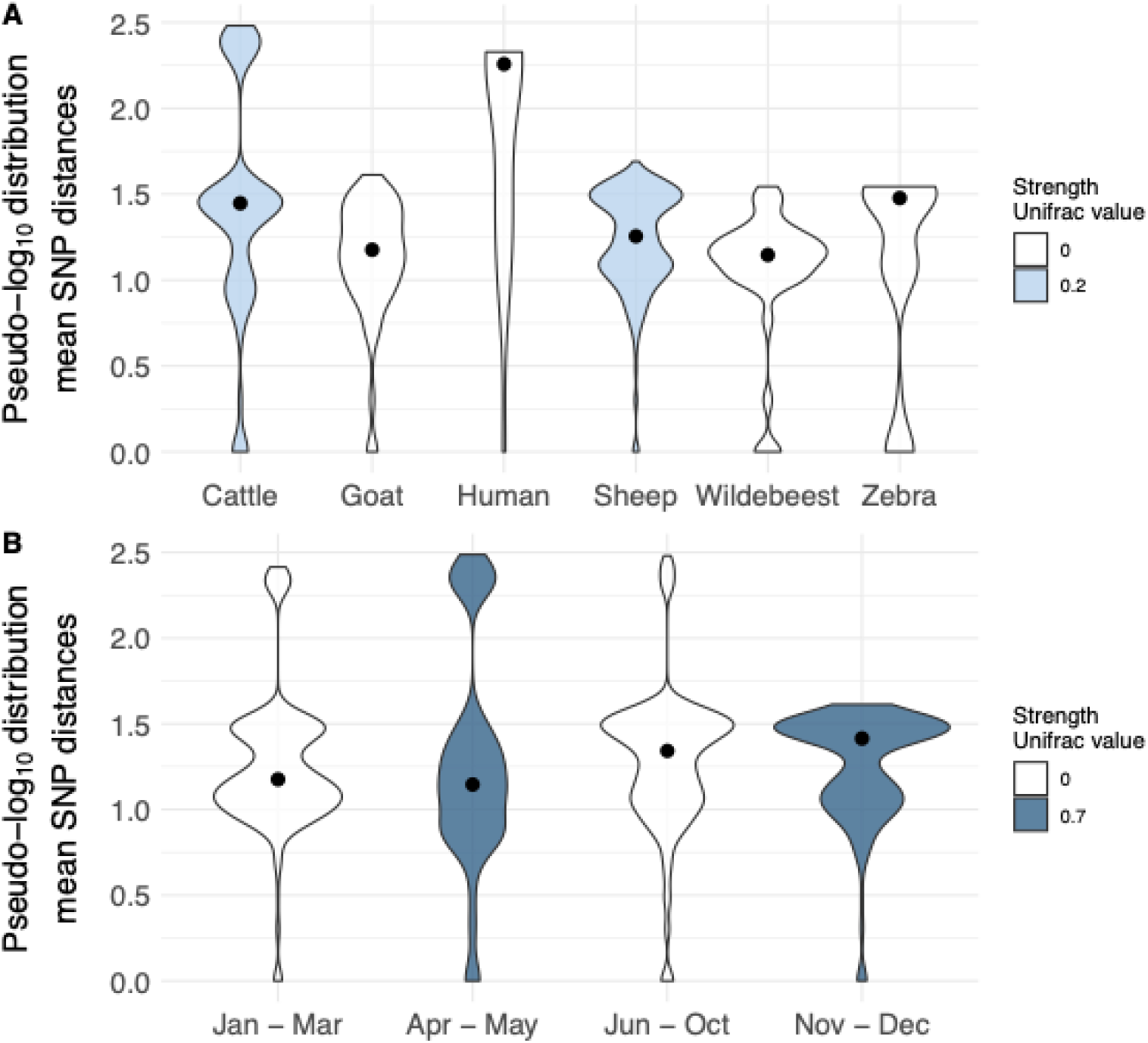
Distribution of mean single nucleotide polymorphism (SNP) distances between Bacillus anthracis sequences grouped by traits and the strength of phylogenetic clustering by trait. Violin plots visualise the distribution of mean SNP distances with a black dot indicating the median value of the distribution. Colour shading represents the strength of the phylogenetic clustering as calculated by the unifrac method. Strength values of 0, represented in white, indicate no stronger clustering signal in the data than observed for a tip-trait permuted control dataset. Values above 0 indicate the n-fold above baseline strength of the unifrac signal. Trait groups with fewer than five sequences were excluded. (A) shows sequences grouped by species and (B) shows sequences grouped by season. A value of 1 was added to all mean SNP distances before logarithmic transformation (pseudo-log_10_) to account for distance values of 0 in the visualisation.

The sequences that had been collected during November through December were sampled from four different years, with the majority of sequences from 2016 and 2017 (16/42 and 19/42, respectively). Spatial distribution of the 29/42 sequences with available GPS data showed the cases were all located in the western region of the NCA, and they broadly structured into two small subregions. These subregions, however, did not correspond with the years, meaning that not all sequences sampled during November and December of a specific year were collected in the same subregion. Sequences collected during April through May were also sampled across four years, with the majority collected in 2017 (15/30). The collection sites for April-May samples are distributed between Endulen and Olbalbal wards (6/30) and are overall less spread out than the samples from November through December. It was not clear why one ELG showed this clustering, as its characteristics, such as number of cases, number of SNP profiles or species affected, did not differ in any evident way from the other ELGs.

## Discussion

In this study, we aimed to characterise the extent of genomic diversity of BA in the anthrax-endemic area of the NCA across different types of hosts – including livestock, wildlife and humans - and different spatial and physical scales. As part of our diversity assessment, we evaluated the use of TC-derived sequences for phylogenetic studies. Secondly, we aimed to test whether factors, such as host or environmental traits, drive BA evolution and population structure.

Our results show that (1) regional clustering is evident, as sequences from the NCA formed a genetically distinct cluster compared to those from neighbouring areas, (2) high genomic diversity exists across the NCA region, within groups of epidemiologically linked cases, individual animal carcasses and different samples from the same animal - with signs of saturation at smaller physical scales, (3) in terms of methodological advances, the TC method enabled us to derive sequences of adequate quality for diversity assessment from non-culturable samples and (4) there was a lack of host specificity of strains that would explain species-specific burden within outbreaks.

### Bacillus anthracis diversity

Previous work suggested that the total diversity of BA in the NCA had mostly been captured (30). However, in this study, we identified a total of 104 SNP profiles, adding 82 additional profiles to the 22 previously described. The sampled area did not increase substantially, but the number of samples of this dataset is almost three times higher than in the previous study, and it includes samples collected from a wider host and temporal range.

We maximised the use of collected material available by implementing a combination of TC and isolate culture methods for sequencing. Our analysis showed that reference-based bait design did not compromise discriminatory power, as previously-defined genotypes were well distinguished by both methods. TC data added an additional nine sequences of previously identified genotypes and enabled us to detect 57 new SNP profiles. In some cases where sequences from the same tissue were compared, TC and isolate derived sequences matched exactly, providing strong evidence that both methods can produce the same results.

We used metadata on collection site and sampling date to assess geographical and temporal distances between cases where identical SNP profiles had been observed. In 18 instances, we observed identical SNP profiles in cases >10 km and at least two weeks apart from one-another, often with more than six months’ separation. The geographic and temporal distance between cases suggests that they arose from separate infection sources, suggestive of widespread genotypes. Due to BA’s ‘time capsule’ effect of extended persistence in soil, long-term circulation and re-occurrence after years – as we observed for the sequences from the wider Arusha region – is actually expected (3, 13). While the circulation of BA types within certain areas has been described for at higher taxonomic levels (i.e. strains from both A and B lineages or MLVA-typing) (28, 75), our study shows that similar patterns are true even for individual SNP profiles.

### The impact of sequence retrieval method on detected diversity

While the implementation of TC for SNP calling does not seem to be associated with any reduction in discriminatory power, the rarefaction analyses of both sequencing methods suggested that TC may introduce some artefactual diversity. Here, extrapolations from TC-derived sequences suggested a much greater underlying diversity than was estimated from isolate-derived sequences. To investigate this effect further, we assessed mean SNP distances among the sequences retrieved by TC in comparison to those retrieved by isolates and we determined the occurrence of gaps (i.e., where there was insufficient read depth to call the allele) and ambiguous sites (i.e., where reads supported more than one possible allele). The mean SNP distances among TC and isolate sequences were very similar, but ambiguous sites were more frequent in TC derived sequences. Since pairwise genomic distance estimation disregards ambiguous and missing sites, one might expect an under- rather than overestimation of diversity. However, our Bali-Phy analyses indicated that such under-estimation was likely negligible within our data.

In agreement with previous findings, we showed evidence for the coexistence of multiple SNP profiles within a single case (3, 30). The higher variation among TC-sequences we observed could thus potentially be due to the underlying metagenomic sequencing approach and subsequent SNP-calling, leading to consensus sequences that are a hybrid of the multiple SNP profiles present. Importantly, we would expect to see ambiguous sites in the assembled reads in these cases. We indeed observed such an ambiguous signal but only in a single case (Supplementary Material 1 Section 1), potentially due to the presence of multiple strains.

In principle TC-derived sequences could provide an opportunity to assess within-sample diversity and to infer the presence of multiple SNP profiles based on ambiguity signal at sites defining previously observed SNP profiles seen in the data set. While we attempted to investigate this option, results were inconclusive (data not shown), likely because much higher read depths than those within our datasets would be required for a robust analysis, as has been shown for viral sequences (76, 77). Another possible reason for higher diversity among TC derived sequences that we can’t exclude would be that the baits captured material of *Bacillus* species other than *anthracis*. Other *Bacillus* species are abundant in the soil (8, 78) and could therefore have contaminated some samples.

In summary, some caution is warranted in calling new SNP profiles detected by TC-derived sequences, since their novelty might be artefactual. However, TC results supported isolate-based findings to a large extent and provided an alternative to culture-based sequencing, thus allowing us to obtain sequence data from several samples where culture was not possible, thereby maximising use of available material.

### Genomic diversity and similarity among and within hosts

A high level of diversity, in terms of the number of SNP profiles detected and their large SNP divergence, was observed between case pairs, within cases, and even within the same sampled material within a host. For the latter, we found maximum SNP distances similar to those seen within-cases overall (28 vs. 32). Due to BA’s low mutation rate (3, 79), it is unlikely that such diversity is generated during the course of one infection (80). This led to our previous hypothesis that the high within-host diversity observed is instead due to a diverse inoculum (30) as has been observed for other bacterial infections (81, 82). We did, however, observe a trend towards lower diversity at smaller physical scales, such as within versus between cases. While this might be partially due to smaller sample sizes, it could also indicate that there is an inherit limit to within-case diversity. This limit might be created by a maximum number of spores entering the host’s body (i.e., typically ingestion/ inhalation for grazing animals and entry through small skin cuts for humans) or competition between strains as they expand within the host. Similar constraints on within-host strain diversity, likely caused by a combination of competition for resources, interaction with the host’s immune system, and specific virulence factors, have been described for other pathogens (81, 82). The factors influencing this are likely system specific, and to our knowledge have been understudied for BA. Extensive sampling from the same carcass, including multiple organs and body sites, would be required to investigate a putative upper limit of within-host BA diversity. Based on our findings, we strongly recommend collecting multiple samples per case whenever possible to adequately capture diversity.

### Factors associated with *Bacillus anthracis* population structure

Interestingly, all 206 sequences from within the NCA clustered within BA lineage 3.2, whereas those obtained from other parts of Arusha and Kilimanjaro Regions fell into lineage 3.1. We also observed that there was much higher genomic homogeneity within the NCA than found among sequences within the Arusha Region (see supplementary Fig S5). Within the NCA, mean SNP distances remained below 50 SNPs, while sequences from the Arusha Region were separated by a few hundred SNPs. This is surprising given that movements of livestock and wildlife in and out of the NCA are likely commonplace, due to the pastoral lifestyle of NCA communities and the lack of physical barriers between protected wildlife areas and neighbouring land. The comparatively lower genomic diversity within the NCA therefore suggests that movement of infected wildlife and livestock from surrounding areas into the NCA might be relatively rare. However, even when considering the NCA only, BA diversity appears strikingly high. Importantly, our increased sampling depth detected higher levels of genomic diversity compared to our previous study (3, 30),with limited signs of saturation on the regional or even case level. This increased diversity is therefore not due to wider geographic sampling, but to denser sampling within the NCA. Wider sequencing from areas surrounding the NCA would be valuable to understand regional patterns of strain circulation.

Our dataset contained multiple groups of samples considered to be potentially epidemiological linked due to their close spatial and temporal proximity (ELGs). We observed that average SNP distances within these ELGs were only slightly smaller than between sequences without such spatio-temporal links. We also still observed extensive strain diversity within a case, involving a large number of phylogenetically divergent sequences. This strongly suggests that despite the appearance of epidemiological links among cases, they may have often originated from multiple different sources. This is not necessarily surprising, as anthrax often causes large scale outbreaks (7, 83), such that many infectious sites (former carcasses) will be located close together and would easily be accessed by an animal grazing in the former outbreak area. Based on the theory that infections are caused by a diverse inoculum, infectious sources themselves might hold a very high diversity of SNP profiles. Both scenarios highlight the extensive environmental soil reservoir of BA spores in this endemic area.

Through the inclusion of TC-derived sequence data we were able to assess for host-association of BA genotypes, comparing sequences obtained from livestock, wildlife and humans. We detected a weak phylogenetic signal in sequences from sheep and cattle, structured into multiple clades. While the statistical test accounts for sample size, the high number of sequences collected from these two host species (n_sheep seq_ = 100 and n_cattle seq_ = 35, respectively) might have provided the critical amount of data to make the signal apparent. It is also possible that livestock movements are more restricted in comparison to those of wildlife, limiting the number of possible sources of exposure. Another possible factor explaining clustering in these hosts is that the NCA comprises a range of eco-climatic zones, with different suitability for particular species. For instance, sheep are increasingly favoured in arid parts of the NCA due to their lower cost compared to cattle and their robustness towards drought (55). Although fewer sequences were available from humans than animals, these were still found throughout the phylogeny and were either identical or closely related to those obtained from livestock, which – in this area – represents the most likely source of infection (i.e., based on more frequent human handling and consumption of livestock than wildlife carcasses). The sampling window for cutaneous human infections was shorter than for an animal that had died, with samples being collected from liquid below the eschar in the former case. Based on our experience, the possible sampling window generally lasted less than seven days (data not shown). Additionally, there is much lower bacterial load in these localised lesions compared with animal carcasses, wherein animals die of septicaemia involving very high levels of circulating bacteria in the blood (approximately 10^8^ colony forming units/ml blood for ruminant species) (19). This makes it comparatively more challenging to obtain sufficient read depth from human versus animal samples during TC. Similarly, wildlife-derived sequences were also phylogenetically diverse, and we found no evidence for lineages or genotypes specific to wildlife or livestock. Seeing certain ruminant host species disproportionately affected during outbreaks compared to others – despite similar susceptibility to BA – is therefore more likely attributable to host ecology-associated factors than strain-associated differences. One limitation of our approach – combining data from isolate and TC sequences – is that only core genome sequence data were available for comparison due to the bait design. BA plasmids contain virulence genes that are both specific and essential to its infectious cycle (84, 85) and plasmid-encoded genes of BA can have host-specific impacts on virulence (42). Since we didn’t systematically obtain plasmid-associated or other accessory gene content within this study, we were unable to assess their potential contribution to differences in host predilection.

When testing for phylogenetic clustering associated with seasons, which differed in terms of their perceived anthrax risk, we found a moderate signal for clustering within both wet seasons (April/May and November/December), in which communities tend to observe fewer cases of anthrax compared with the two dry seasons. Lower case numbers could conceivably arise from a smaller number of spore reservoirs, resulting in higher similarity between strains (86). But there may also be a link to promoted spore formation under wet conditions (49), which could allow spores from a carcass to become available again within the same season in which the animal died. We did not see SNP profiles re-occurring during the same seasons across multiple years, meaning that the observed clustering is constrained to the same year and could thus be tied to areas that provided either particularly attractive or infectious grazing sites at the time based on the climatic conditions of the given year (14, 87). According to studies in other parts of Africa, seasonality of case occurrence can correlate with species. Interestingly, with our dataset, wildebeest were only documented, and thus sampled, during January to March, the dry season – while reports for Kruger and Etosha National parks describe the peak of anthrax deaths of this species during the wet seasons (52). In our dataset, livestock cases were recorded fairly evenly throughout the year, but with a peak during June to October (the long dry season).

We also tested for phylogenetic clustering in association with a previously-described high-risk infection zone identified through participatory mapping with the NCA communities (46). Sequences collected within this high-risk area were separated by on-average shorter SNP distances, while sequences collected outside of the high-risk zone were separated by larger SNP distances. The reasons for this remain speculative. We obtained 58 sequences from within, in contrast to 155 sequences from outside the high-risk zone, which is likely connected to the fact that the high-risk zone covers a smaller portion of the full sampling area. Beyond the likely explanation that a larger area may contain more diverse strains, pastoralists typically take their livestock on daily grazing routes within <10 km of their settlements (46). This suggests that infections and deaths are more likely to occur in proximity to settlements, with the accumulation of similar strains, especially when considering that settlements will more likely be established in areas with proximity to favourable grazing sites. At the time of the analysis, we did not have access to the locations of settlements in the area, particularly those that are non-permanent or semi-permanent, to evaluate whether their location could drive the differences in SNP differences observed between the low- and high-risk areas.

## Conclusion

Using a combination of isolate and targeted-capture (TC)-derived sequences, we were able to conduct a unique investigation into BA diversity at the One Health interface in an anthrax endemic area, as well as into factors that shape its population structure. While we observed intuitive patterns of shorter genetic distances among cases occurring in close spatio-temporal proximity, as well as sequences from the same case, extensive diversity of BA was present at all levels, and multiple unrelated genotypes were often identified in clusters of anthrax cases that appeared likely to be epidemiologically linked. Our broad dataset allowed us to test for phylogenetic clustering based on host species and other spatial and environmental variables. We detected only a weak clustering signal in BA sequences from sheep and cattle and found no evidence for species-specific lineages. Instead, identical or very closely related sequences were shared among livestock, wildlife and humans. Our results suggest that host behavioural differences such as time spent in certain habitats and grazing areas along with herd aggregations, are more likely to explain why some species appear disproportionately impacted during anthrax outbreaks.

## Author Contributions

Conceptualisation: T.L.F., T.L., R.B., A.H. Data curation: A.H., M.M., T.L.F. Formal analysis: A.H., M.M., T.L.F. Funding acquisition: T.L.F., B.T.M., T.L., R.B., S.J.L, A.H., H.O.A., S.N. Investigation: D.M., T.L.F., T.L., A.H., S.K.M., B.W. Methodology: A.H., M.M., B.W., S.J.L., R.B., T.L.F. Project administration: T.L.F., B.T.M., T.L., I.K. Resources: I.K., Z.E.M., H.O.A., S.N., M.P.R., J.A.C. Supervision: R.B., T.L., S.J.L., T.L.F. Visualisation: A.H., T.L.F. Writing – original draft: A.H., T.L.F. Writing review and editing: all authors.

## Acknowledgements

We are grateful for the support received from the NCA community and authorities. We thank the Ngorongoro District Council, Ngorongoro Conservation Area Authority, District Veterinary Office, Tanzania Wildlife Research Institute and members of our field team Sabore Ole Moko, Kadogo Lerimba, Allutu Masokoto, Mary Lonyori, Christopher Kiboya, and Nicholaus Sitayo – for assistance with this study. We also thank the Directorate of Veterinary Services, Ministry of Agriculture, Livestock and Fisheries, and Ministry of Health for their support. In addition to funding from the MFA of Finland, we also acknowledge the Finnish Centre for Military Medicine (SOTLK) for its support through the project “Strengthening Biosafety and Biosecurity in Tanzania by Biodetection Capacity Building” (2014–2024), jointly implemented with the TVLA. We thank Marco van Zwetselaar for bioinformatics assistance. We also thank Venance Maro from the KCMC-Duke University collaboration, as well as Alex R. Mremi and Patrick T. Amsi from the Department of Pathology, Kilimanjaro Christian Medical Centre, Moshi.

## Supporting Information

Supplementary Material 1. Text and Figures.

Tables S2. Unifrac Results

Supplementary Material 2. Microreact Readme.

